# Mutation severity spectrum of rare alleles in the human genome is predictive of disease type

**DOI:** 10.1101/835462

**Authors:** Jimin Pei, Lisa Kinch, Nick V. Grishin

## Abstract

The human genome harbors a variety of genetic variations. Single-nucleotide changes that alter amino acids in protein-coding regions are one of the major causes of human phenotypic variation and diseases. These single-amino acid variations (SAVs) are routinely found in whole genome and exome sequencing. Evaluating the functional impact of such genomic alterations is crucial for diagnosis of genetic disorders. We developed DeepSAV, a deep-learning convolutional neural network to differentiate disease-causing and benign SAVs based on a variety of protein sequence, structural and functional properties. Our method outperforms most stand-alone programs and has similar predictive power as some of the best available. We transformed DeepSAV scores of rare SAVs observed in the general population into a mutation severity measure of protein-coding genes. This measure reflects a gene’s tolerance to deleterious missense mutations and serves as a useful tool to study gene-disease associations. Genes implicated in cancer, autism, and viral interaction are found by this measure as intolerant to mutations, while genes associated with a number of other diseases are scored as tolerant. Among known disease-associated genes, those that are mutation-intolerant are likely to function in development and signal transduction pathways, while those that are mutation-tolerant tend to encode metabolic and mitochondrial proteins.

## Introduction

Genetic variations are major determinants of human diseases and phenotypes [1]. Accelerating pace of large-scale sequencing projects on genomes and exomes has greatly expanded the landscape of human genetic variations. It remains a challenging task to assess the functional impact of these variations [2]. Comprehensive analysis of genetic variations, especially those found in and near the exons of protein-coding genes [3], may shed light on gene-disease relationships and provide insight into the mechanisms of diseases and variations in phenotypes [4]. The increasing number of sequenced human genomes and exomes from the general population would enhance the statistical power of such analyses [5].

Different types of genetic variations occur at varying scales ranging from large structural variations such as chromosomal rearrangements and copy number variations (CNVs), to insertions and deletions (indels) of up to hundreds of nucleotide positions, and to single-base-pair (single-nucleotide) variations (SNVs) [6]. Any types of genetic variations could cause human diseases with a variety of mechanisms, including effects on chromatin organization, gene expression and regulation, protein function, and genetic instability [7-11]. The observed frequencies of genetic variations in the general population are tied to their fitness cost as well as the evolutionary history of the human species and its ancestors. While common variations, most notably SNVs, were first documented, more rare genetic variations (e.g., those with minor allele frequency (MAF) less than 0.0001) at the individual level have been identified in large-scale sequencing projects of the general population [5] as well as patients with certain diseases such as cancer [12] and intellectual disability [13]. Although some recurring variations have been identified to be the drivers of diseases, a significant number of rare mutations are persistently found, and their clinical significance are difficult to evaluate. Genome-wide association studies can pinpoint to the genetic loci, mostly marked by common SNVs, that exhibit statistically significant associations with diseases or phenotypes [14, 15]. Association of rare and de novo mutations to common and rare diseases could be unveiled through familial or trio studies that are facilitated by genome or exome sequencing nowadays [16, 17]. Coupled with pathway profiling, systematic analysis of genetic variations in patients could shed light on the biological processes underlying diseases [18]. However, disease gene prioritization and disease-causing variation discovery are still difficult [19, 20].

The identity change in a single base pair position is the most common type of genetic variation. In protein-coding regions, non-synonymous variations (missense mutations) result in the change of a single amino acid in the protein product [21]. Clinical consequences of these missense mutations, referred to as single amino acid variations (SAVs), are generally more difficult to evaluate than synonymous mutations (generally benign) and nonsense (stop codon) mutations (often resulting in loss of function). A number of computational methods [22] have been developed to assess the mutational effects of SAVs found in the human proteome encoded by around 20,000 protein-coding genes.

Essential genes compromise the viability of an individual when their function is lost. Such genes can be identified by observing intolerance to loss-of-function variants at the population level [23]. In genetic terms, essential genes tend to exhibit haploinsufficiency, where the loss of one of two gene alleles is detrimental. Genetic alterations of haploinsufficient genes are not only a major cause of dominant diseases [24], but also play key roles in developmental disorders [17]. On the one hand, haploinsufficient genes can function as tumor supressors [25]. On the other hand, essential genes tend to be expressed at higher levels in cancer cells than in normal cells [26]. Thus, knowledge about gene essentiality can help prioritize deleterious variants in genetic studies and could help prioritize therapeutic targets in cancer. Given the role of essential genes in human disease, considerable efforts have gone into developing methods for haploinsufficiency prediction [5, 27-30].

In this study, we developed a deep convolutional neural network-based method for predicting the clinical impact of human SAVs based on analysis of sequence, structural and functional properties of SAVs in the human proteome. The neural network prediction results of SAVs observed in the general population were used to calculate a mutation severity measure that estimates tolerance of each human protein-coding gene to deleterious missense mutations. This measure correlates with gene essentiality and specific disease classes such as cancer and autism. Finally, we observed a dichotomy of mutation severity for disease-associated genes: those that are mutation-intolerant tend to function in development and signal transduction pathways, while those that are mutation-tolerant tend to function in metabolism.

## Results and discussion

### Analysis of human disease-related genes and their variants

We obtained a set of likely pathogenic (disease-causing) genetic variants from two database resources: ClinVar [31] and UniProt [32]. ClinVar aggregates reported variant-disease associations from submissions of research studies. ClinVar variants annotated as “Pathogenic” or “Likely pathogenic” were found in ∼4,200 protein-coding genes, about one fifth of the human proteome. SAVs were found in the majority (3,410) of these genes. Non-SAV variants were also found in most of them (∼3,300 genes). Non-SAV variants include indel variants, single-nucleotide variations in noncoding regions (mostly at splice sites), nonsense single-nucleotide variations (to stop codons), and a small number of synonymous variants (Figure 1A). The 31,171 SAVs made up about 30% of all ClinVar variants (Figure 1B). UniProt is another curated resource for likely pathogenic SAVs. The number of proteins with UniProt SAVs annotated as disease-related is 2,755 (Figure 1C). Most of these genes (2,590) overlap with the ClinVar disease-associated gene set, with UniProt contributing only 165 disease-associated genes not found in the ClinVar set. On the other hand, more than half of the UniProt pathogenic variants (15,697 out of 29,300, Figure 1D) were not found in the set of ClinVar pathogenic variants. The total number of likely pathogenic variants in the unified ClinVar and UniProt set is ∼47,000. We also obtained a set of benign variants (∼45,000) by combining the ClinVar variants annotated as “Benign” or “Likely benign” and the UniProt variants in the category of “Polymorphism”.

**Figure 1.**
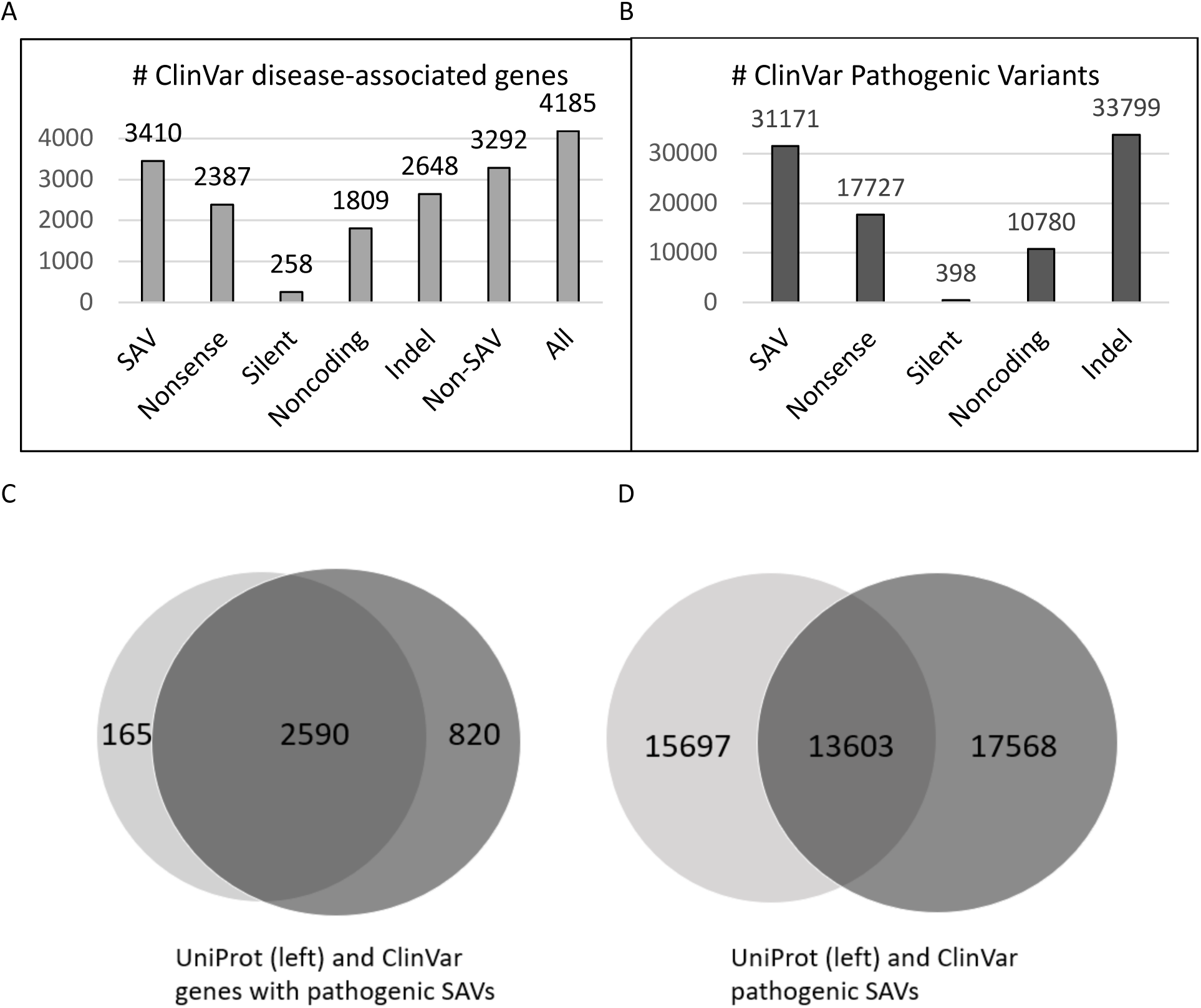
Distribution of disease-associated genes and variants. **A)** The number of ClinVar disease-associated genes with different types of variants. The non-SAV category combines the categories of nonsense, silent (synonymous), noncoding, and indel. **B)** The number of variants of different types in ClinVar disease-associated genes. **C)** Venn diagram of genes with pathogenic SAVs from UniProt and ClinVar. **D)**. Venn diagram of pathogenic variants from UniProt and ClinVar.

The number of likely pathogenic SAVs are not evenly distributed among the disease-associated genes. The three genes with the greatest number of SAVs encode long proteins: FBN1 (Fibrillin-1, 2,871 amino acids), LDLR (Low-density lipoprotein receptor, 860 amino acids), and SCN1A (Sodium channel protein type 1 subunit alpha, 2,009 amino acids), each of which has more than 500 pathogenic SAVs. In part, it may be due to the length of these proteins. 75 genes possess more than 100 pathogenic SAVs. More than half of the disease-associated genes with SAVs (2,003 out of 3,575) have less than 5 pathogenic SAVs, and 916 of them have only one pathogenic SAV. One cause of the uneven distribution of SAVs could be the bias in research studies of common diseases and certain genes (e.g., the *LDLR* gene involved in hypercholesterolemia).

### Enrichment analysis of sequence, structure and functional properties in likely pathogenic SAVs and gnomAD SAVs

We compiled a set of protein sequence, structure, and functional properties (features) predicted by computer programs or retrieved from UniProt Feature fields (see Materials and methods). A log-odds score was used to determine if any feature is enriched or depleted in amino acid positions with pathogenic SAVs compared to the background frequency of that feature in all human proteins (see Materials and methods). We observed a 1.7-fold enrichment of conserved positions (Consv3 in Figure 2A) and more than 3-fold depletion of variable positions (Consv1 in Figure 2A) in pathogenic SAVs. Similarly, results of three disorder prediction programs (DISOPRED3 [33], SPOT-Disorder [34], and IUPred2A [35]) consistently show that ordered regions are enriched and disorder regions are depleted in pathogenic SAVs. Predicted β-strands and α-helices are slightly preferred in pathogenic SAVs, while coil regions of secondary structure prediction, low complexity regions, and coiled coil regions are disfavored.

**Figure 2.**
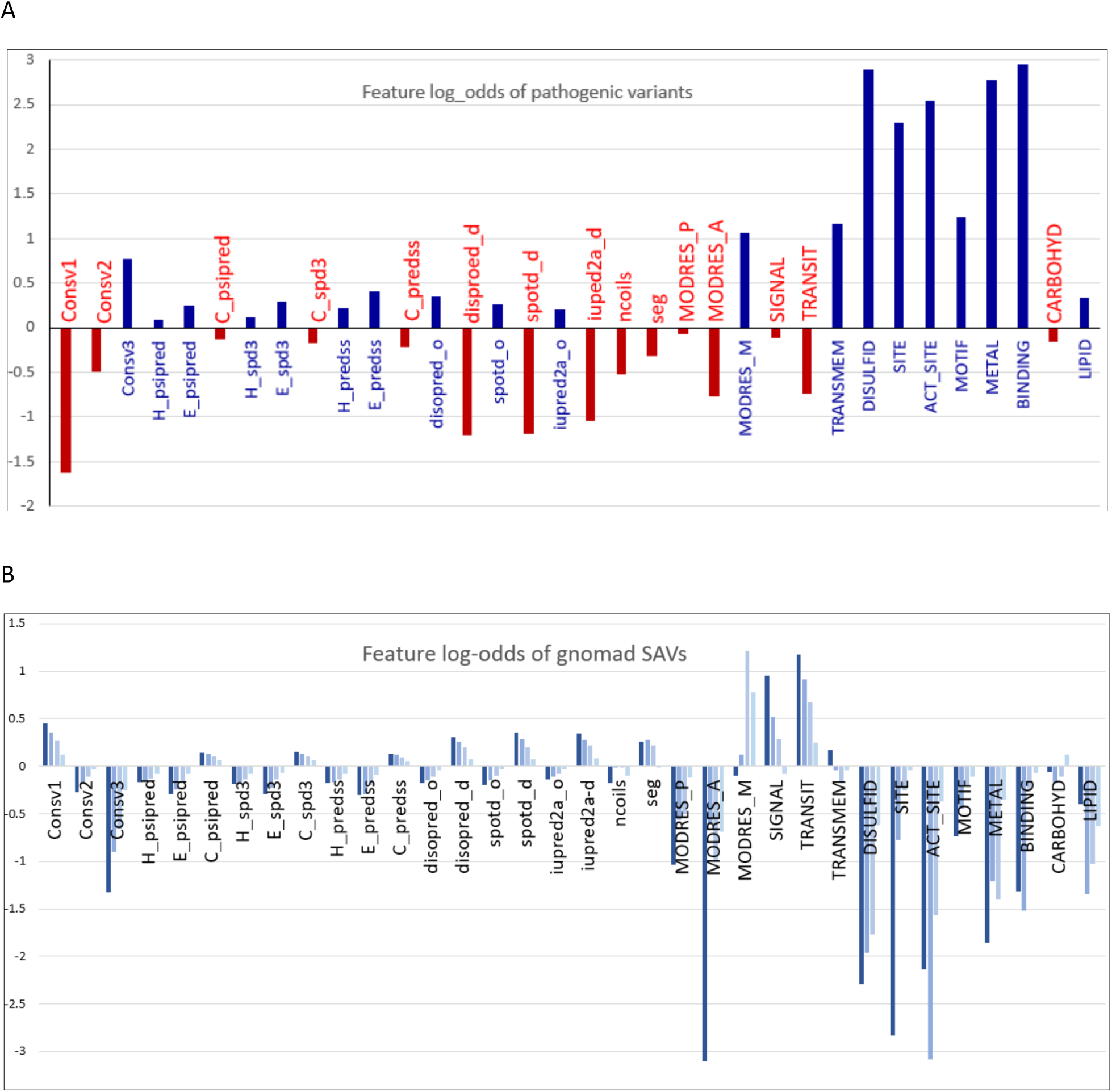
Enrichment of SAVs among sequence, structure and functional properties. **A)** Enrichment/depletion of features in pathogenic SAVs (y-axis shows log2 based log-odds scores). **B)** Enrichment/depletion of features in gnomAD SAVs with different MAF ranges (from light blue to dark blue: MAF < 0.0001, 0.0001 ≤ MAF < 0.001, 0.001 ≤ MAF < 0.01, 0.01 ≤ MAF).

For regions with indications of subcellular localization, signal peptides and mitochondrial transit peptides are depleted in pathogenic SAVs, but transmembrane segments are enriched by more than 2 fold. Several UniProt features showing the strongest enrichments in pathogenic SAVs are related to protein stability (UniProt feature DISULFID: cysteine residues participating in disulfide bonds) or function (UniProt features: SITE, ACT_SITE, METAL, MOTIF, and BINDING, see their explanations in Materials and methods). Except the MOTIF feature, they exhibit more than 4-fold enrichment in pathogenic SAVs (log2-based odds score more than 2, Figure 2A).

We also analyzed SAVs found in more than 12,000 exomes (>24,000 alleles) in the gnomAD [5] database, which provides a comprehensive catalogue of natural variants from the general population. Common SAVs (MAF ≥ 0.01) should be mostly benign, and they only make up a small fraction of gnomAD SAVs (about 0.5%). The gnomAD database possesses many more rare SAVs with MAF less than 0.01, a significant portion of which are singletons (found only once in all exomes). We partition gnomAD SAVs according to their MAFs into four categories (MAF < 0.0001, 0.0001 ≤ MAF < 0.001, 0.001 ≤ MAF < 0.01, and MAF ≥ 0.01). Enrichments of protein sequence, structure, and functional features in each gnomAD SAV category were analyzed in the same way as for pathogenic SAVs (Figure 2B). Common gnomAD SAVs (MAF ≥ 0.01) generally exhibit opposite enrichment/depletion trends compared to pathogenic SAVs. Features such as DISULFID, SITE, ACT_SITE, METAL, MOTIF, and BINDING exhibit the most prominent depletion in common gnomAD SAVs and the strongest enrichment in pathogenic SAVs. In contrast, features enriched in common SAVs include variable positions (Consv1), coil regions of secondary structure prediction, predicted disordered regions, low complexity regions, signal peptides, and mitochondrial transit peptides. The enrichment or depletion of features were gradually curtailed when moving from the category of common gnomAD SAVs to less frequent gnomAD SAV categories (Figure 2B). This behavior suggests that many low frequency SAVs, especially those with MAF less than 0.0001 in the general population could be deleterious, because functionally important residues (specified by UniProt features SITE, ACT_SITE, BINDING, METAL, and MOTIF) are found more frequently in these rare SAVs than in the common SAVs.

### DeepSAV – a deep neural network-based method for SAV pathogenicity prediction

We developed an artificial intelligence method (DeepSAV) that uses a deep-learning convolutional neural network to predict SAV pathogenicity based on input features of sequence, structure, and functional information (see Materials and methods). The features include amino acid type, sequence profile, sequence conservation, secondary structure and disorder predictions, coiled coil and low complexity region predictions, sequence regions indicating subcellular localization (signal peptide, transit peptide, transmembrane segments), and functional and stability properties from the UniProt database such as post-translational modifications, disulfide bond, active site, and motifs. Features of a window of 21 amino acid positions centered around the mutated amino acid were encoded as input. The neural network has mainly convolutional layers and applies techniques such as max-pooling, residual network, and dropout (supplemental Figure S1). It is trained on a large set (40,000 pathogenic and 40,000 benign) of SAVs from the ClinVar and UniProt database.

Testing on an independent set of SAVs (3,000 pathogenic and 3,000 benign), DeepSAV yields better performance (measured by area under the ROC (receiver operating characteristic) curve (AUC)) to differentiate pathogenic from benign SAVs than many stand-alone programs such as SIFT [36], polyphen-2 [37], PROVEAN [38], CADD [39], LRT [40], MutPred [41], MutationAssessor [42], PrimateAI [43], and a simple baseline fitness score we used before in the Critical Assessment of Genome Interpretation (CAGI) evaluations [44] (Figure 3A). DeepSAV’s performance is also better than two meta-predictors MetaSVM [45], MetaLR [45] that use predictions results of a number of other stand-alone predictors (Figure 3A). DeepSAV trails the two best methods REVEL [46] (a meta-predictor) and VEST4 by about 0.01 in AUC. We were not able to find information about the algorithm of VEST4, an improved version of VEST [47], which shows the best performance, and used VEST4 scores as given in dbNSFP [22].

**Figure 3.**
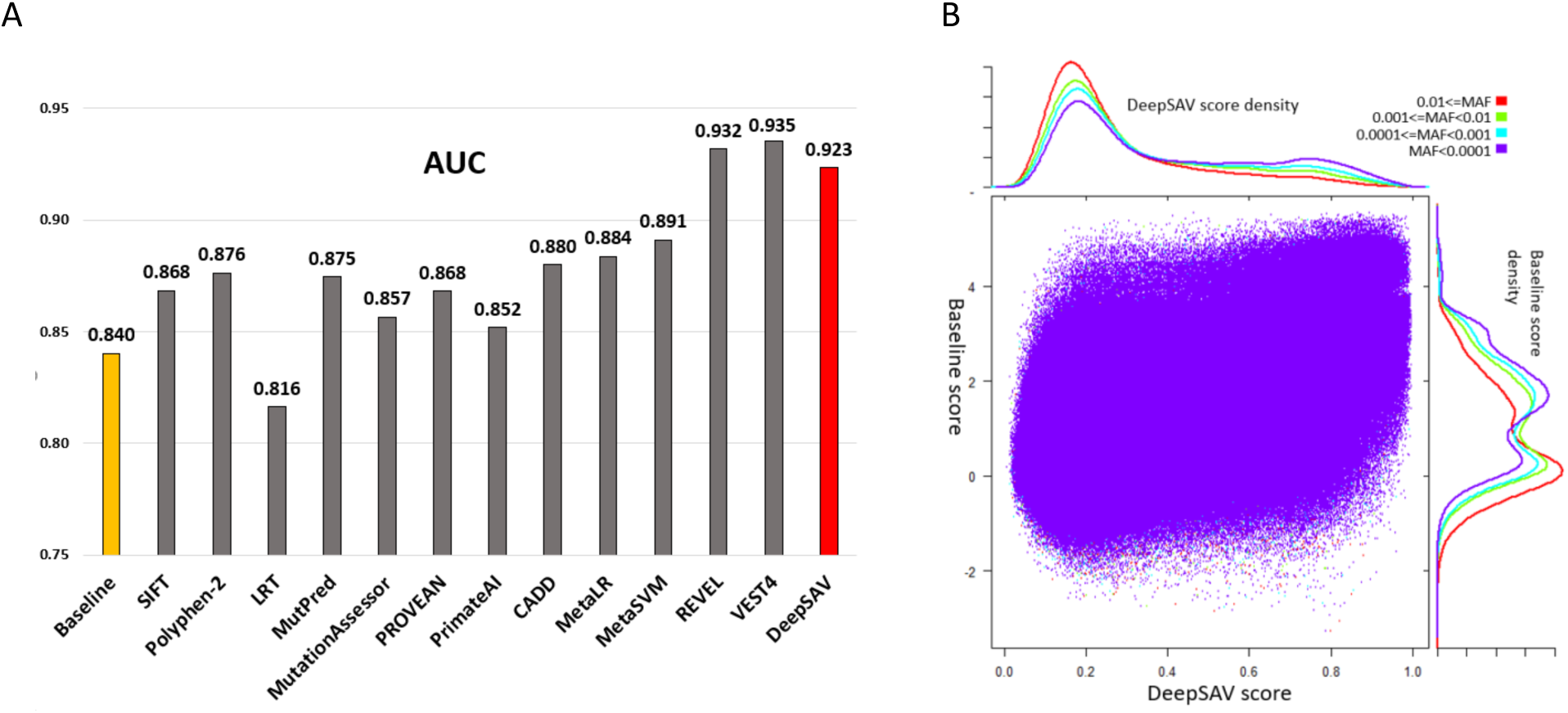
**A)** Performance of variant pathogenicity prediction programs in terms of AUC (area under the ROC curve) measure. **B)** Scatter plot of DeepSAV scores and baseline fitness scores for SAVs observed in gnomAD. Datapoints for four different MAF categories are shown. Their density plots are shown by the axes above (DeepSAV) and right (baseline fitness).

We calculated DeepSAV scores and baseline fitness scores for human protein SAVs observed in the gnomAD database. They show a positive correlation (correlation coefficient: 0.57, Figure 3B). They both exhibit bimodal distributions for SAVs in each of the four different MAF categories (MAF ≥ 0.01 (common SAVs), 0.001 ≤ MAF < 0.01, 0.0001 ≤ MAF < 0.001, and MAF < 0.0001). The range of DeepSAV scores is between 0 and 1, with higher scores suggesting increasing likelihood of being deleterious (pathogenic). For common SAVs (MAF > 0.01), the distribution of DeepSAV exhibits a high peak in the low score range, and a flat tail in the high score range, suggesting that the majority of common SAVs are predicted to be benign. With increasing stringencies of rare SAVs, the volume of the peak in the low-score range decreases while the tail in the high-score range increases, suggesting that pathogenic SAVs are more likely to occur in rarer SAVs. The baseline fitness scores display similar behavior for SAVs in different MAF categories, although the peaks in high and low scoring ranges appears to overlap more compared to the DeepSAV scores.

### Mutation severity scores enrich for essential genes with potential disease associations

Deep sequencing of human exomes has highlighted the contribution of rare SAVs to gene function and complex diseases [48, 49]. Certain genes may be more tolerant to deleterious or partially deleterious SAVs due to their functional properties. To evaluate the mutation tolerance of genes, we transformed our DeepSAV predictions of SAVs present in the human population (from the gnomAD database [5]) into an average mutation severity measure for each gene (AvgAI scores, see Materials and methods). AvgAI scores based on our deep neural network predictor were calculated for SAVs with several filters for common variants (MAF less than 1, 0.01, 0.001, or 0.0001), and were compared to the same scores calculated using a simple baseline predictor [44] (AvgBF) (see Materials and methods). For rare SAVs (MAF < 0.0001) the baseline AvgBF score correlates well 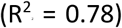 with the AvgAI score (Figure 4A), suggesting that the deep neural network predictor reflects the profile score difference between amino acids of major and minor alleles that go into the baseline predictor.

**Figure 4.**
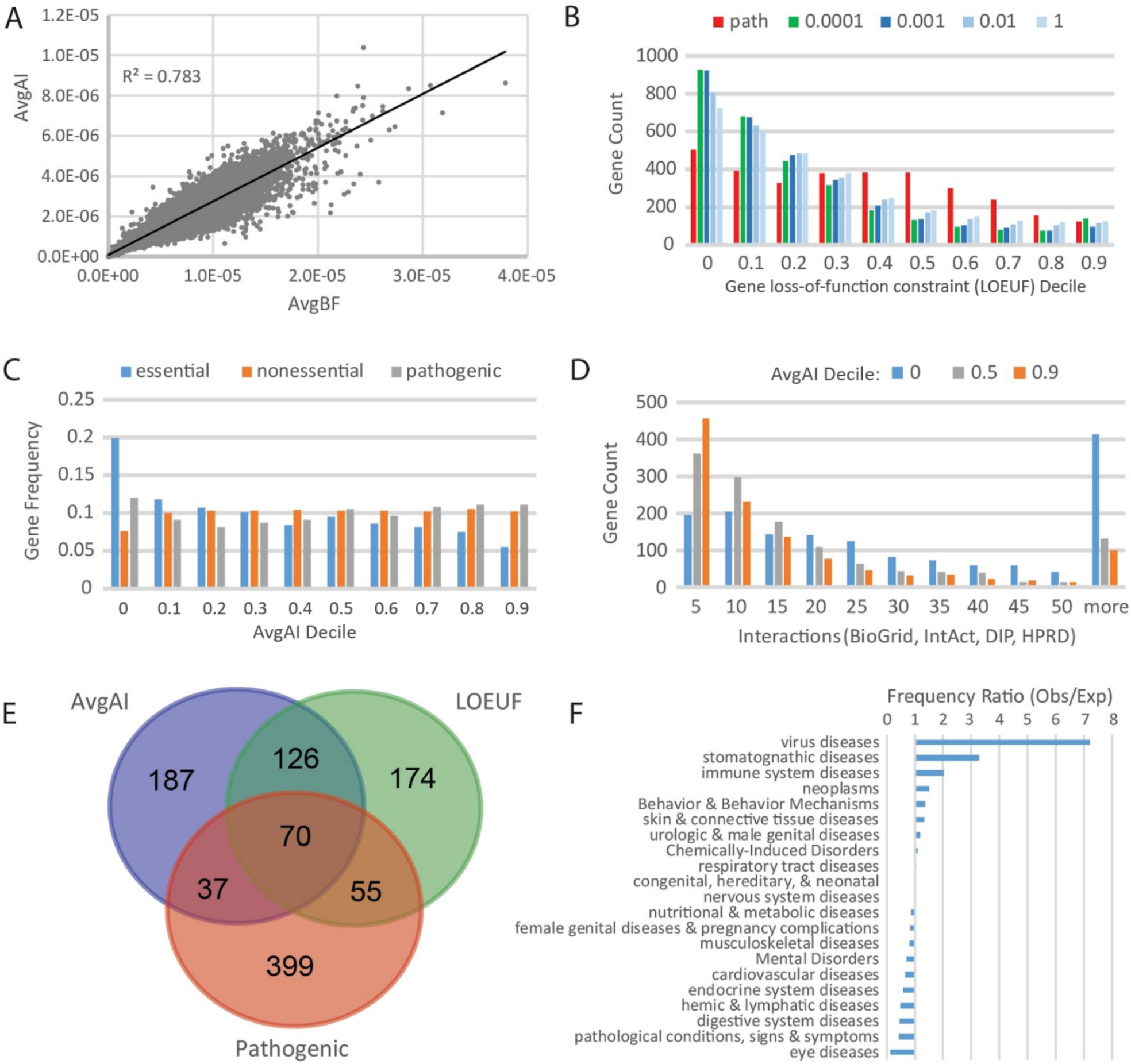
Mutation severity measures based on DeepSAV identify potential disease-associated genes. **A)** Mutation severity measure based on DeepSAV scores (AvgAI) correlates with the measure based on baseline fitness scores (AvgBF) for 17,480 human genes **B)** Distribution of gene count among decile bins of loss-of-function constraint measure (LOEUF) for a set of genes (>3,000) with pathogenic SAVs (red bars, labeled as “path”) and for the gene sets (having the same number genes) with the lowest AvgAI scores computed at various cutoffs of minor allele frequencies (0.0001, 0.001, 0.01 and 1). On the x axis, 0 means the first LOEUF decile [0, 0.1] (the same extrapolation applies to other numbers). **C)** Distribution of gene frequency among AvgAI deciles (MAF cutoff 0.0001) for the same gene set with known pathogenic SAVs compared to essential and nonessential gene sets. **D)** Distribution of protein interactions from four databases (BioGrid [50], IntAct [51], DIP [52] and HPRD [53]) integrated in PICKLE [54] for gene sets within three different mutation severity AvgAI score deciles (0, 0.5 and 0.9). **E)** Venn diagram highlights overlap among essential genes with known pathogenic variants (labeled as “Pathogenic”), essential genes with lowest loss-of-function constraint scores (LOEUF), and essential genes with lowest mutation severity measure (AvgAI). **F)** Representation of disease class associated with genes from the overlapping set of top-ranked genes by LOEUF and AvgAI (126 genes, not including genes with known pathogenic SAVs).

Human genes have been classified by a measure (LOEUF) that reflects their tolerance to inactivation (loss-of-function) [5]. To see how our AvgAI score correlates with the LOEUF score, we ranked human genes from low AvgAI (mutation-intolerant) to high AvgAI (mutation-tolerant). The LOEUF distribution for top-ranking mutation-intolerant genes was compared to that of a set of known disease-associated genes (Figure 4B). The top-ranking mutation-intolerant genes selected by the mutation severity measure (lowest AvgAI scores) include progressively more loss-of-function constrained genes with increased filtering of common variants. In contrast, the disease-associated gene set displays a bimodal distribution of highly constrained genes at low LOEUF and less constrained genes at median LOEUF. Thus, the mutation severity measure for rare SAVs reiterates a gene’s tolerance to inactivation, with top-ranking mutation-intolerant genes being more frequent in the percentile of lowest tolerance to inactivation.

A fraction of the disease-associated human gene set (17%) is annotated as essential by one or more CRISPR screens [55, 56]. Among all curated gene-disease associations in DisGeNET (May 2019 version) [57], the essential disease-associated genes contribute to 2,477 diseases or syndromes and 1,847 neoplastic processes. We originally reasoned that genes able to accumulate numerous detrimental SAVs (evaluated by high AvgAI scores) were less likely to contribute to disease phenotypes. However, the AvgAI scores do not discriminate disease-associated genes collectively, giving similar gene frequencies displayed across the AvgAI deciles (Figure 4C, gray bars). Instead, AvgAI scores tend to select for gene essentiality, with an increase in essential genes and a decrease in non-essential genes at the lowest AvgAI decile (Figure 4C). Similar to noted trends of both essential and disease-associated genes [58, 59], human genes with AvgAI scores in the lowest decile, regardless of their essentiality, exhibit increased numbers of protein interactions than those from higher AvgAI deciles (Figure 4D).

Although the loss-of-function constraint measure LOEUF and the mutation severity measure AvgAI display similar trends in reflecting gene essentiality, they define different gene sets that might be used to evaluate potential new disease-associated genes. A comparison of essential genes with the lowest LOEUF scores, essential genes with the lowest AvgAI scores, and essential genes with pathogenic SAVs highlights the divide among these gene sets (Figure 4E). The overlap between low-AvgAI set and low-LOEUF set (126 genes, not including genes with pathogenic SAVs) provides a potential source of disease-associated genes. Indeed, despite the lack of documented pathogenic SAVs in the 126 mutation- and inactivation-intolerant genes, curated DisGeNET gene-disease associations annotate almost half (55 genes) as being involved in disease. The set includes almost all disease classes, with several being over-represented when compared to all gene-disease associations: including virus diseases, stomatognathic diseases, immune system diseases, and neoplasms (Figure 4F). Given their propensity to associate with disease, the essential genes selected by our AvgAI measure could provide insight into novel gene-disease associations.

### Mutation-intolerant essential genes cluster with disease-associated genes and contribute to diseases

The essential genes with pathogenic SAVs (70 genes, Figure 4E) that overlap with both the loss-of-function-constrained genes (lowest LOEUF) and the low mutation severity genes (lowest AvgAI) set a standard to prioritize other potential disease-associated genes (126 overlapping genes shared by the low LOEUF and the low AvgAI sets, but without pathogenic SAVs, Figure 4E). Clustering these two sets of genes (70+126) together using complete linkage of correlated distances over six scores (see Materials and methods) places potential disease-associated genes among those that are known to be associated with diseases (Supplemental Figure S2). Two clusters (40 genes) with the highest proportions of disease-associated genes exhibit lower AvgAI scores than other clusters, indicating their intolerance to detrimental missense mutations (red labels in Supplemental Figure S2). Inspection of gene-disease associations for genes in these two clusters reveals that 68% are linked to curated diseases.

Enriched GO biological process terms are similar for each identified gene cluster, and annotation clustering of terms from the combined set (40 genes from two clusters, 17 with pathogenic SAVs) highlights their function in RNA splicing (enrichment score 9.02, 13 genes), gene expression (enrichment score 6.81, 31 genes), and chromosome segregation (enrichment score 4.34, 9 genes). Two disease-associated genes and eleven others belong to the most enriched cluster and function in RNA splicing, including pre-mRNA processing factor 3 (*PRPF3*) having variants associated with Retinitis pigmentosa and splicing factor 3b subunit 1 (*SF3B1*) having variants associated with acute myeloid leukemia, among other neoplastic processes. Five of the potential disease-associated genes involved in RNA splicing (*DHX15, HNRNPH1, SRSF1, PCBP2*, and *DHX9*) are reported to be associated with myelodysplasias in DisGeNET [57], and the spliceosome has become a therapeutic target for myeloid malignancies [60, 61].

The third most enriched functional cluster includes six disease-associated genes and three others that function in chromosome segregation. Three of the disease-associated genes (*RAD21, SMC3*, and *SMC1A*) functioning in chromosome segregation have genetic variants causing Cornelia de Lange syndrome (CdLS), which manifests developmentally as intellectual and growth retardations. The protein-coding products of these genes comprise three of the four subunits of the mitotic cohesion complex responsible for chromosome segregation. Mutations in this complex are known to cause a number of diseases termed cohesinopathies, of which CdLS is the best characterized [62]. One additional chromosome segregation gene from this set, PDS5 cohesin associated factor A (*PDS5A*), is associated with CdLS in DisGeNET literature [63]. Gene dosage appears to be an important component of CdLS severity, which is consistent with the essential nature of our selected gene set [23].

### Mutation-intolerant disease-associated genes function in development and signaling pathways

Over half of the top 1,000 human genes ranked by low AvgAI (571 genes) are associated with 1,618 diseases, 262 phenotypes and 184 disease groups such as “Intellectual Disability” that encompass multiple similar diseases or phenotypes. To understand the functional context of mutation-intolerant genes that are associated with disease, we assigned them to pathways in Reactome [64] (467 genes). Functional enrichment of these pathways highlight involvement in axon guidance (P-value < 1.78E-15), development (P-value < 9.64E-14), and neurotransmitter receptors and postsynaptic signal transmission (P-value < 1.12E-13), among others. Those genes in the enriched category of “developmental biology” describe early steps in development that give rise to diverse tissues in the body and thus represent critical processes that should contribute to fitness. In fact, 25% of this gene set participate in development, and many are annotated as essential (47 genes) or conditionally essential (17 genes). However, a significant portion of the developmental genes are not considered essential (52 genes). Many of them encode protein kinases (18 genes), homeobox transcription regulators (4 genes) or proteins with other signaling domains that are expanded in the genome like rho-binding domains (3 genes), pleckstrin homology domains (4 genes), or SH3 domains (3 genes).

While a relatively small core set of essential genes exists in eukaryotes whose loss of function results in lethality, a larger subset of genes exhibits conditional lethality that also affects fitness [65]. For example, deleterious mutation of immune system genes might not necessarily result in a lethal phenotype. However, their contribution to survival under specific conditions like being challenged with an infectious agent could be considered as essential. This spectrum of gene essentiality is indeed reflected in the disease-associated genes functioning in development, as they exhibit essential and conditionally essential responses in CRISPR screens [55, 56]. Furthermore, many of the mutation-intolerant and disease-associated genes not considered as essential belong to families like protein kinases that have expanded in the human genome and could be functionally redundant [66]. Thus, the concept of gene essentiality alone does not necessarily suggest the ability to cause disease.

The mitogen-activated protein kinases ERK1 and ERK2 function in development and signal transduction pathways. They represent a duplication that is thought to be functionally redundant [67]. However, ERK2 includes two known pathogenic variants that are associated with various neoplastic diseases (E322K) as well as with inborn genetic disease (R135T). The ERK2 structure (Figure 5A) includes a relatively small set (9 positions) of DeepSAV-predicted deleterious SAVs from the gnomAD database (DeepSAV score >0.75). One of these SAVs (D106G) lines the ATP-binding pocket, and four are buried in the structure core (D44Y, G136E, R148H, and R194T), with R148 belonging to the HRD motif that controls kinase activation. The rest are in a C-terminal extension to the catalytic domain that lines the surface of the kinase in between the N-lobe and the C-lobe. The known pathogenic variants cluster together with many of the predicted deleterious mutations. Thus, while this kinase is thought to be functionally redundant, some variants have been reported as pathogenic, several others are predicted as detrimental, and the gene is intolerant to deleterious mutation (AvgAI score 3.24E-7 and ranked 142 out of more than 17,000 genes). Accordingly, the ERK2 gene was shown to be conditionally essential in a CRISPR screen [55], suggesting conditions exist where the functional redundancy of the two kinases breaks down.

**Figure 5.**
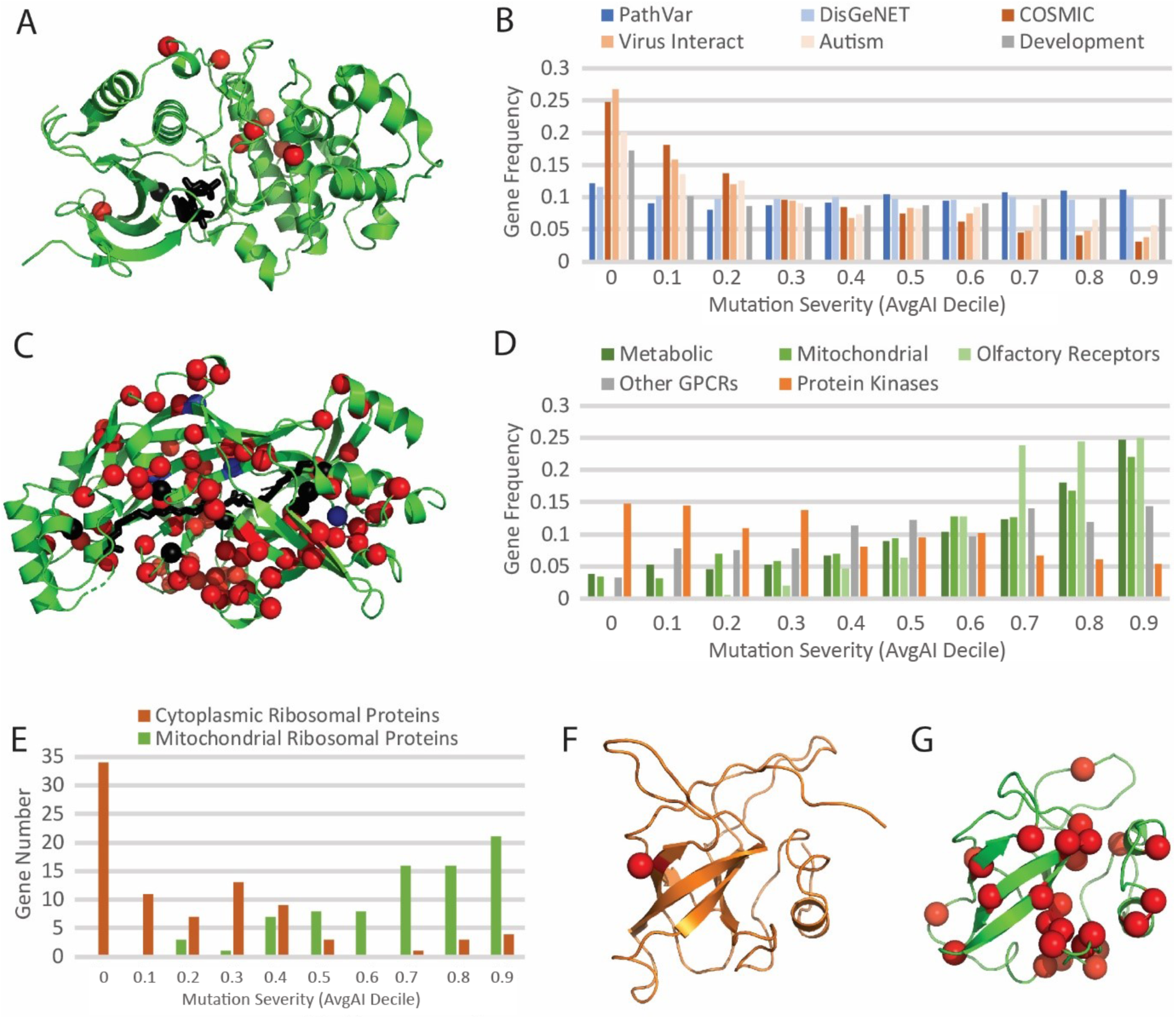
Mutation-intolerant and mutation-tolerant genes prefer different pathways and disease types. **A)** Top ranked AvgAI genes like ERK2 kinase (PDB 4fmq) have relatively few DeepSAV predicted deleterious variant positions (DeepSAV score > 0.75, red spheres). One of these (black sphere) is near (< 4Å) the active site (ANP substrate analog in black stick). **B)** Mutation severity spectrum of disease-associated genes measured by their frequencies in AvgAI deciles. Associated disease for each gene set is labeled above. **C)** Bottom ranked AvgAI genes like CD36 (PDB 5lgd) are tolerant to predicted deleterious mutations (DeepSAV score > 0.75, red spheres), including several positions (black spheres) lining the fatty acid (black stick) binding sites or with known pathogenic variation in platelet glycoprotein deficiency (blue spheres). **D)** Mutation severity spectrum of pathway gene sets and large paralogous gene sets measured by their frequencies in AvgAI deciles. **E)** Mutation severity spectrum of ribosomal proteins functioning in the cytoplasm (orange bars) and in the mitochondria (green bars) measured by their numbers in AvgAI deciles. **F)** 60S ribosomal protein L23 from cytoplasm (Ribosomal protein L14P homolog, PDB: 6ek0, chain LV) in orange cartoon has a single detrimental predicted SAV (red sphere). **G)** Mitochondrial 39S ribosomal protein L14 (PDB 5oom, chain L) in green cartoon has multiple predicted detrimental SAVs.

Mutation intolerance appears to be a quality exhibited not only by genes associated with developmental disorders [17], but also by genes contributing to other various disease types such as cancer (COSMIC) [12], autism [68] (https://gene.sfari.org), and viral interacting proteins [69] (Figure 5B). However, mutation severity does not select for collective disease-associated gene sets (PathVar (ClinVar and UniProt genes with pathogenic SAVs) and DisGeNET, Figure 5B), with nearly uniform distributions of the number of genes among AvgAI deciles. Such a preference for mutation-intolerance in selected disease types suggests that the AvgAI score can be particularly useful for prioritizing disease genes when combined with additional considerations, such a disease type or functional pathways contributing to the disease state.

### Mutation-tolerant genes function in metabolic pathways and mitochondria

The concept of functional redundancy from gene duplication extends not only to critical components of developmental and signal transduction pathways, but also to those of metabolic pathways [59]. Enriched functional pathways of mutation-intolerant genes that are associated with disease highlight repeated involvement of core genetic information processing (e.g., transcription and RNA processing) and signal transduction components, but they tend to exclude those of metabolism. In fact, mutation-tolerant genes (a numeric matched set of genes with the highest AvgAI scores) are significantly enriched in metabolism (P-value < 2.05E-13) in the Reactome pathway database [64].

An example of a mutation-tolerant gene product is platelet glycoprotein 4 (CD36), which functions in cell adhesion by serving as a receptor for thrombospondin in platelets as well as in the metabolism of lipids through binding long chain fatty acids. CD36 represents one of the most mutation-tolerant genes in the diseases-associated set, with 155 DeepSAV-predicted detrimental mutations (DeepSAV score>0.75) in 98 positions covering the structure, including seven lining the fatty acid binding site (Figure 5C). Although this gene is tolerant to mutation, known pathogenic variants (I413L, R386W, P90S, and F254L), with I413L lining the fatty acid binding pocket, cause platelet glycoprotein deficiency, a congenital disease of the hemic and lymphatic class. Furthermore, DisGeNET associates this gene with metabolic phenotypes of impaired glucose tolerance, insulin resistance, and insulin sensitivity. While mutations in CD36 can still lead to disease, the mutation tolerance of the gene might be explained by the recessive nature of the associated disease, by the ability of two paralogs, SCARB1 and SCARB2, to serve as functional replacements, or by the tissue-specific nature of the disease [70].

Therefore, a dichotomy seems to exist for disease-associated genes, where those that are mutation-intolerant tend to function in development and signal transduction pathways, while those that are mutation-tolerant tend to function in metabolism. These trends imply a greater overall fitness cost of mutations in developmental and signal transduction genes when compared to metabolic genes. However, extreme functional redundancy in some signal transduction proteins may lead to their tolerance to mutations. The mutation severity spectrum of signal transduction proteins with numerous paralogs that could exhibit functional redundancy are shown in Figure 5D. Paralogous olfactory receptors (OR), which represent a specialized set of G protein-coupled receptors (GPCRs) that detect odors, are more mutation-tolerant than other GPCRs. In fact, human OR paralogs include more pseudogenes [71] (464, not included in Figure 5D) that have accumulated enough mutations to render them inactive than functional genes (361, included in Figure 5D), and this well-known OR variability likely contributes to an individual’s sense of smell [72]. Both non-OR GPCRs (Figure 5D, gray bars) and protein kinases (Figure 5D, orange bars) shift in the spectrum towards mutation intolerance when compared to either metabolic enzymes [73] (Figure 5D, dark green bars) or nucleus-encoded proteins functioning in the mitochondria [74] (Figure 5D, medium green bars), organelles that provide energy from nutrients using metabolic processes [75].

Metabolic enzymes exhibit similar tendency towards mutation-tolerance as the ORs (Figure 5D). One explanation for the greater tolerance of metabolic genes to mutations might be the redundancy not only in gene duplications, but also in non-homologous proteins that can serve as functional analogs of the same reactions [76]. Metabolites can also be acquired through transport mechanisms, relieving the evolutionary constraints on certain metabolic enzymes. Finally, metabolic pathways exhibit both redundancy and plasticity, allowing for multiple ways to arrive at the same metabolite [77].

The mutation-tolerance observed for nucleus-encoded mitochondrial proteins might reflect their roles in metabolic processes [75]. However, this tendency is also exhibited by the ribosomal proteins that function in mitochondria compared to ribosomal proteins functioning in cytoplasm (Figure 5E): the majority of mitochondrial ribosomal proteins have high AvgAI scores while the majority of cytoplasmic ribosomal proteins have low AvgAI scores. As an example, side-by-side comparison of ribosomal L14P/L23E-like proteins functioning in the cytoplasm (L23, Figure 5F) and the mitochondria (mitoL14, Figure 5G) highlights the mutation-intolerance and mutation-tolerance, respectively. Both proteins adopt similar small 5-stranded meander barrel folds with relatively long loops that interact with RNA in the assembled ribosome. However, the cytoplasmic L23 includes only a single predicted pathogenic variant (I40F, DeepSAV score = 0.788), while the mitoL14 includes 34 predicted pathogenic variants (all but one are rare with MAF < 0.0001) covering 28 positions in the 145 residue-long protein. Neither of these examples possess known pathogenic variants, and only the cytoplasmic version is associated with a neoplastic process in DisGeNET. This marked difference in rare allele mutation severity cannot be explained by either domain or pathway redundancy. The main function of mitochondria is to supply energy, which can be partly salvaged by increasing nutrient intake and decreasing energy-demanding activities. In addition, mitochondria might be able to overcome lowered fitness of mutations in the ribosome through their processes of fusion and fission that help maintain both functional properties and the integrity of the mitochondrial genome that harbors the mitochondrial ribosomal RNA genes [75].

### Mutation-intolerant and mutation-tolerant genes function in different disease classes

The set of mutation-intolerant genes define several over-represented disease classes, including virus diseases, behavior & behavior mechanisms, stomatognathic diseases, hemic & lymphatic diseases, immune system diseases, musculoskeletal diseases, nervous system diseases, neoplasms, pathological conditions, signs & symptoms, and mental disorders (Figure 6A). Development and signal transduction are enriched among the mutation-intolerant genes associated with these specific disease classes. Furthermore, top mutation-intolerant genes tend to participate in relevant functional pathways. For example, mutation-intolerant genes associated with behavior diseases are enriched in neurotransmitter receptors and postsynaptic signal transmission (P-value < 1.11E-16), and those involved in immune system diseases are enriched in cytokine signaling of the immune system (P-value < 1.4E-14).

**Figure 6.**
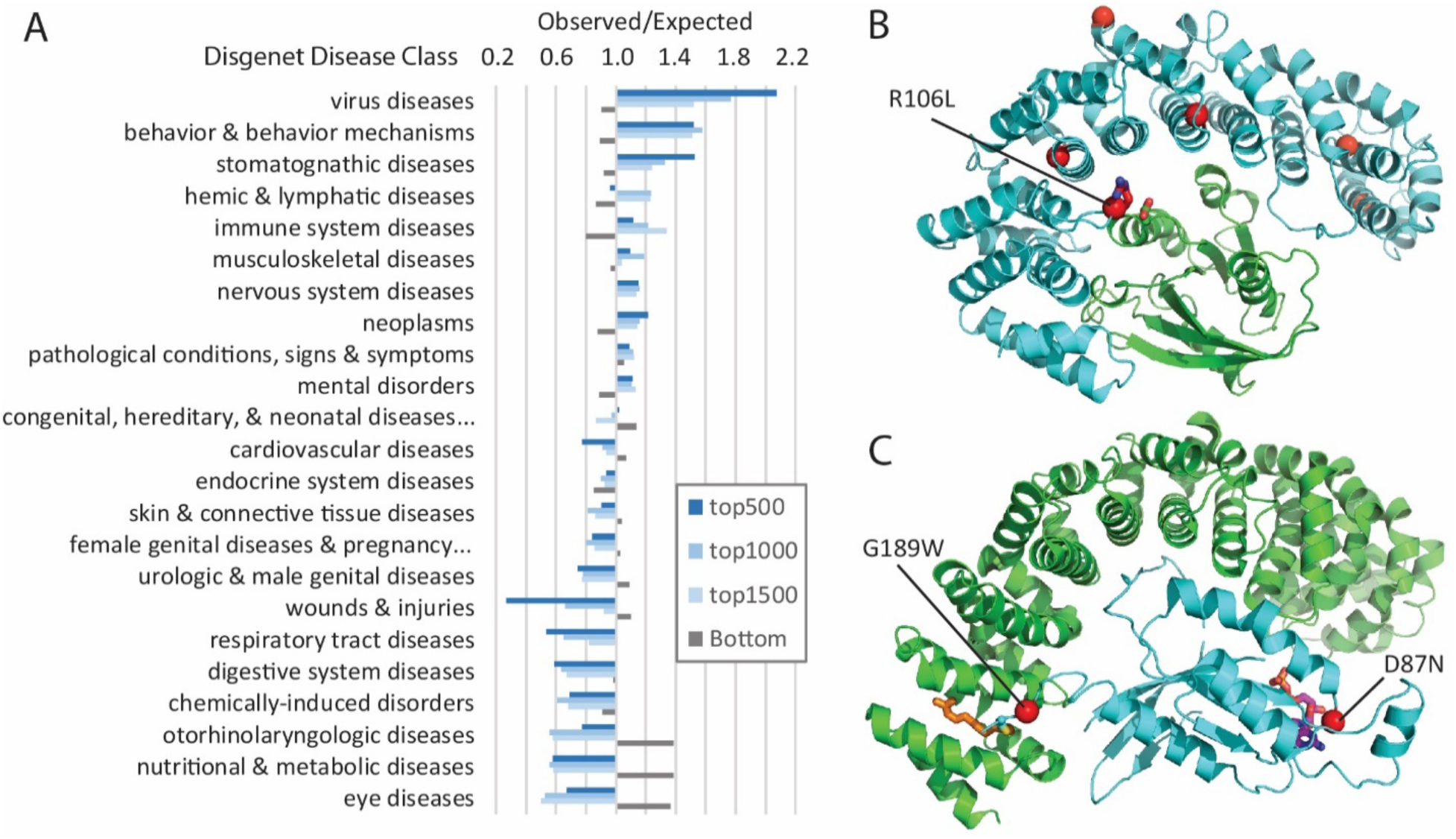
Mutation-intolerant genes exhibit pathway preferences and are exploited by viruses. **A)** Genes are ranked from low to high by mutation severity measure, AvgAI. The top ranked genes are mutation-intolerant, and the bottom ranked are mutation-tolerant. Ratios of observed/expected frequencies of disease class associations for sets of mutation-intolerant (top-) and mutation-tolerant (Bottom) genes are shown. Diseases are ordered by the exp/obs frequency ratios in the top1000 set (top 1000 genes with the lowest AvgAI score). **B)** Ribbon diagram of KPNB1 (cyan) bound to Ran GTPase (green) with DeepSAV-predicted detrimental variants (red spheres), including R106L (stick) at the interaction interface (from PDB 1ibr). **C)** Ribbon diagram of GEF (green) bound to RHOA GTPase (cyan, PDB 5zhx), with labeled DeepSAV-predicted detrimental variants (red spheres) adjacent to a farnesylation site (orange stick) and near the active site (stick colored by atom, from superimposed GTPase 1tx4).

The disease classes that are the most under-represented in mutation-intolerant (and over-represented in mutation-tolerant) genes include eye diseases, nutritional and metabolic diseases, otorhinolaryngologic diseases, chemically induced disorders, digestive system diseases, respiratory tract diseases, and wounds and injuries. These disease classes tend to be either tissue-specific or related to metabolism. For example, eye diseases involve genes functioning in visual perception, in cilium morphogenesis, as structural constituents of the eye lens, and in phototransduction. Alternatively, nutritional diseases involve genes of respiratory electron transport, steroid metabolism, TCA cycle, and fatty acid metabolism, among others. The nutritional diseases associated with mutation-intolerant genes tend to be dominated by a clinically heterogeneous group of disorders that arise as a result of dysfunction of the mitochondrial respiratory chain (mitochondrial diseases), as well as by obesity and diabetes that display a range of severity in affected individuals and can develop in adolescence or later in life. The relatively modest impact of these diseases on survival may be a reason for the genes associated with such diseases to tolerate mutations.

### Viruses exploit disease-causing mutation-intolerant genes for infection

Viral diseases are the highest over-represented disease class among mutation-intolerant genes. Potentially, viral strategies for successful replication and evasion of host immunity could benefit from targeting essential genes that accumulate fewer mutations. In fact, similar observations of viral proteins interacting with more evolutionarily constrained host genes suggest that viruses have driven close to 30% of adaptive amino acid changes in the human proteome, with HIV infection causing a statistically significant increase in adaptation [69]. These evolutionarily constrained viral-interacting host proteins tend to be mutation-intolerant (Figure 5B), while a set of similarly highly adaptive proteins that interact with *Plasmodium* [78] do not have the same degree of preference for mutation-intolerance (data not shown).

Over a third of the virus disease gene set is involved in HIV coinfection, which describes simultaneous infection of a single host cell by two or more virus particles. Identified HIV coinfection-associated gene products function in pathways such as signaling by interleukins/cytokines, regulation of RUNX3, stabilization of P53, and host interactions with HIV factors, among others (ordered by Reactome enrichment). Cytokines, including interleukins, play a critical role in immunity. Because HIV infects immune CD4 T cells, the connection to interleukin/cytokine signaling molecules that regulate T cell growth and differentiation (i.e. through IL2 or CCL2) is known [79, 80], and two of the interleukin signaling examples (PSME3 and PSMC3) represent biomarkers for the disorder [81]. A significant portion of the HIV-related genes (19 out of 23 or 82%) are annotated as essential, including all host interaction factors (*KPNB1, RAN, PSMC3, PSME3, PSMA5, PSMA6, PSMC5, and PSMA4*), supporting the notion that infection strategies involving essential proteins are utilized by HIV, and potentially other viruses.

The essential HIV host factor importin subunit beta-1 (KPNB1) includes several gnomAD SAV positions predicted as detrimental using both DeepSAV and baseline fitness scores. However, the gene does not belong to our disease-causing set and has no disease associations in DisGeNET. KPNB1 mediates nuclear import of ribosomal proteins [82], and also works together with the RAN GTP-binding protein to bind and import HIV Rev into the nucleus where it exports viral mRNAs for translation [83]. In a structure of KPNB1 bound to RAN (Figure 6B) [84], these positions are buried (T150P, L238S, and A389V) or partially buried (R234G, C436Y) in the hydrophobic core of the KPNB1 repetitive α-hairpins. Such variations could result in local structure instability and loss of function. One surface SAV, R105L, interacts with a nearby E in RAN. Replacement of R by L, which removes a potential interaction of charges, could lower the key interaction of KPNB1 with RAN that drives nuclear import of HIV Rev.

Another example of a GTP-binding protein, RHOA, contributes to viral diseases such as Burkitt Lymphoma, which is a cancer of the lymphatic system with a subtype linked to Epstein-Barr virus (EBV) [85]. RHOA variants are deemed likely pathogenic for several other neoplastic disorders, and several missense variants are listed in DisGeNET, although not in association with Burkitt Lymphoma. Despite the apparent tumor-promoting effects of RHOA in various cancers, previous studies suggest mutations of the gene in the case of Burkitt lymohoma and other neoplastic processes are inhibitory [86, 87]. There are only two predicted detrimental SAVs in RHOA in gnomAD. One of them (G189W) maps to the disordered C-terminus adjacent to a residue that gets farnesylated. The disordered and modified C-terminus adopts a coil structure when bound to the RAP1GDS1 guanine nucleotide exchange factor (GEF) (Figure 6C), and the replacement of a small G to W with the larger sidechain would incur steric problems in the GEF-bound conformation. Similarly, a larger sidechain adjacent to the farnesylation site might reduce the modification and influence RHOA localization. While the second SAV (D87N) is relatively conservative, its position near the GTP binding pocket adjacent to a K sidechain that mediates Guanine nucleotide binding might influence enzymatic activity.

## Materials and methods

### Human proteome, sequence alignment, and baseline fitness score

The human proteome was obtained from the UniProt database (version 2018.12) [32]. The orthologous groups of human proteins were obtained by OrthoFinder [88] applied to a set of representative vertebrate proteomes. For human proteins in large orthologous groups, we replaced their orthologous groups by the ones retrieved from the OMA database [89] that are usually much smaller and thus exhibit better alignment quality. Multiple sequence alignments of orthologs were obtained by MAFFT [90]. Sequence profile of each position of an alignment, represented as the estimated amino acid frequencies, was calculated as described before [91]. For any amino acid change, we used a previously devised baseline fitness score to represent the severity of the mutation, based on the log-odds ratio between original amino acid and mutated amino acid [44, 92].

### Positional features used in impact predictions of deep convolutional neural network

For each human protein position, we deduced features reflecting amino acid type, sequence profile, sequence conservation, structure properties, and available functional annotations. The type of 20 amino acids is used as one feature with one-hot encoding. Both the original amino acid and the variant amino acid are encoded in this way, resulting in 40 features. The estimated amino acid frequencies of each position in the multiple sequence alignment of orthologs were used as 20 features. Sequence conservation scores of the multiple sequence alignment of orthologs were calculated by AL2CO [93] and used as one feature. Prediction of 3-state secondary structures (helix, strand, and coil) were made by three programs (PSIPRED [94], SPIDER [95], and PSSpred [96]), resulting in nine features. Three features are based on disorder propensities predicted by three programs (DISOPRED3 [33], SPOT-Disorder [34], and IUPred2A [35]). In addition, low complexity region predictions by SEG [97] and coiled coil predictions by NCOILS [98] were encoded as two features. We also used features reflecting protein-targeting or functional regions or positions from the UniProt sequence annotations. Regions of N-terminal signal peptide (indication of proteins going through secretory pathway), transit peptide (indication of mitochondrion targeting), and transmembrane segments were obtained from UniProt feature records SIGNAL, TRANSIT, and TRANSMEM, respectively. Three post-translational modifications (phosphorylation, acetylation, and methylation) were extracted from the UniProt MODRES records. Other UniProt Features includes DISULFID (cysteines participating in disulfide bonds), CARBOHYD (site with covalently attached glycan group), METAL (binding site for a metal ion), BINDING (binding site for any chemical group (co-enzyme, prosthetic group, etc.)), ACT_SITE (amino acid directly involved in the activity of an enzyme), SITE (any single amino acid site that could be functionally relevant), LIPID (site with covalently attached lipid group(s)), and MOTIF (short, i.e. up to 20 amino acids, sequence motif of biological interest). For 1-dimensional convolutional network, the above 89 features from a window of 21 amino acids (the target position and 10 neighboring positions on each side) were used as input. Features in neighboring positions beyond the first or last residues were zero-filled (zero-padding). One additional feature encodes the indicator of zero-padding for such positions (1 for positions beyond the first or last residues, and zero for normal amino acid positions within the protein length). The number of features for each position is 90. By using a window of 21 positions, a total of 90 × 21=1890 values serve as the input of the convolutional neural network for each training and testing data point.

### Architecture and hyperparameters of the deep-learning convolutional neural network

We used a deep-learning artificial neural network for prediction of SAV pathogenicity. The diagram of neural network structure is shown in supplemental Figure S1. It consists of seven 1-dimensional convolutional (conv1d) layers, two max-pooling layers, and two dense layers before the output. The residual network architecture is implemented twice by combining the input of a conv1d layer with the output after several layers of that input (thick arrows, supplemental Figure S1). The initial input has a window size 21 and 90 channels corresponding to 90 features encoding protein sequence, structure and functional properties (described above). The number of filters and the kernel size of other conv1d layer are 200 and 3, respectively. Each of the two dense layers has 100 nodes and has a following dropout layer with the dropout rate of 0.5. The ReLU activation function is used in all layers except the output layer that uses the softmax function. The batch size is set to 128 in the training process. The neural network was written in python with the TensorFlow package. The prediction score of any SAV, ranging from zero to one, reflects the likelihood of the SAV being pathogenic, and is termed DeepSAV score.

### Training and testing dataset for neural network

We obtained SAVs that were classified as likely pathogenic from the ClinVar [31] and UniProt database. For the ClinVar database, these SAVs are classified as “Pathogenic” or “Likely Pathogenic”. For the UniProt database, these SAVs are classified as “Disease” in the SwissVar database [99]. Benign SAVs are those classified as “Benign” or “Likely Benign” in the Clinvar database and those classified as “Polymorphism” by SwissVar in the UniProt database. The non-redundant sets of pathogenic and benign SAVs were randomized and 40,000 of each category are used for training. During the training process, a small subset of these SAVs (1,000 pathogenic and 1,000 benign) were used as a testing set to monitor the performance after each epoch of training using the remaining training SAVs. The training was stopped after the decrease of performance over this small testing set to prevent overfitting. For evaluations of the performance of the deep neural network in this study and other predictors (scores obtained from the dbNSFP database [22]), 3,000 independent pathogenic and 3,000 independent benign cases were set aside from the training SAVs and used as the testing dataset.

### Enrichment analysis of features in likely pathogenic SAVs and gnomAD SAVs

he enrichment score is defined as the logarithm of the ratio between two probabilities. This ratio is the probability of observing a feature among a subset of amino acid positions (e.g., positions with pathogenic SAVs, or positions with gnomAD SAVs with MAF in a certain range) divided by the probability of observing that feature among all amino acid positions in the human proteome. It reflects enrichment (if the log-odds score is above zero) or depletion (log-odds score less than zero) of the feature in the subset compared to the background (the whole proteome).

### Quantification of mutation severity of gnomAD SAVs at the gene level

We applied our deep neural network method to the prediction of mutational impact of gnomAD [5] SAVs obtained from the dbNSFP database [22] (version 4.0). Rare SAVs were defined as those with MAF less than a certain cutoff (e.g., 0.01, 0.001, 0.0001). For any given MAF cutoff, the cumulative mutation severity measure according to our predictions is calculated as

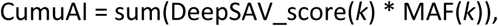

where DeepSAV_score(*k*) is the DeepSAV score of any rare SAV *k*, and MAF(*k*) is its minor allele frequency. The average mutation severity score of rare SAVs is defined as

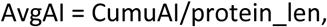

where the normalization factor is the protein length (protein_len) of the gene. Similarly, the average mutation severity measure according to baseline fitness predictions is calculated as

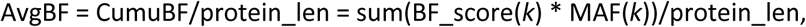

where BF_score(*k*) is the baseline fitness score of any rare SAV *k* in the gene.

### Analysis of mutation severity measures for potential disease-associated genes

Average mutation severity scores calculated using baseline fitness (AvgBF) [92] or our deep neural network (AvgAI) predictors were transformed into percentiles (using Excel percentrank) for 17,480 human protein-coding genes. For comparison of our average mutation severity scores to constrained genes that are more likely to be detrimental when inactivated (LOEUF score [5]), we transformed LOEUF scores by percentrank (for 16,670 human genes with LOEUF score). The resulting gene count distributions among LOEUF deciles were plotted for a set of genes with pathogenic SAVs or the top 3,284 genes ranked by four sets of AvgAI scores from lowest (unlikely to acquire damaging mutations) to highest (tolerates acquired mutations). We chose the AvgAI mutation severity measure filtered at MAF<0.0001 for further evaluation and transformed the score into decile rank. For the disease-related gene set, we compared decile rank of AvgAI score to those of human genes annotated as essential by either of two large-scale CRISPR experiments [55, 56] (2,108 genes) or annotated as non-essential (11,589 genes) in both.

To identify potential disease-associated genes, we compared essential genes having the lowest AvgAI and LOEUF scores with essential genes with pathogenic SAVs using a venn diagram. The overlap between the AvgAI and LOEUF sets is considered to be enriched for potential disease-associated genes. The overlapping set (126 genes) was assigned to disease classes using the DisGeNET curated gene-disease associations (GDAs) [57]. We removed group and phenotype associations from the GDAs. MeSH (Medical Subject Headings) disease class frequencies for the set (observed frequencies) were compared to disease class frequencies assigned to all curated genes (expected frequencies) to evaluate over- and under-representation (observed/expected frequency ratios).

To further select among potential disease-related genes, we clustered the gene set (126 genes) together with the genes with pathogenic SAVs (70 genes) using ClustVis [100] with six measures for each gene (AvgAI, LOEUF [5], lnGDI [29], number of rare gnomAD mutations (MAF filter: 0.0001), HIPred [27], and P(HI) [24]). The raw scores for each measure were converted to *Z*-scores and were pre-processed with row centering and no scaling. Principal component analysis using the SVD with imputation option indicated the first and second components explain 40.8 % and 25.9% of the data variance, respectively. Scores were plotted as a heatmap from high (red) to low (blue) *Z*-score, and both genes and measures were clustered using complete linkage of correlation distances. The genes were split into three large groups for visualization (supplemental Figure S2), with the top 20 clusters separated by space in the resulting heatmaps. Functional analysis for potential disease-related genes were performed using DAVID clustering (medium stringency with 0.001 ease) of GOfat biological process terms [101] and GO enrichment of PANTHER classification [102].

### DisGeNET mapping

We mapped all human genes with AvgAI scores to curated DisGeNET diseases (using UniProt to GeneID provided in the Downloads section from the DisGeNET website [57]). Of the 9,414 total GeneIDs with curated GDAs, we mapped AvgAI scores and ranks from the complete AvgAI dataset to 8,426 genes with curated GDAs. Over-representation (enrichment) and under-representation (depletion) of disease classes from MeSH were calculated over various sets of genes as the ratio of the observed frequency of each class to the expected frequency of each class calculated from disease class frequencies in the entire curated gene-disease database. We chose sets of genes for plotting the distributions of overrepresented disease classes over all AvgAI ranks, where genes ranked up to 1,000 (top1000 set) tend to include increased frequencies, and genes ranked higher than 12,000 (bottom set) tend to have stable frequencies. We included additional sets surrounding the top1000 (top500 and top1500) to observe trends. We excluded disease classes with few representatives, including F02: psychological phenomena & processes, C22: animal diseases, C03: parasitic diseases, C01: bacterial infections & mycoses, C24: occupational diseases, and C21: disorders of environmental origin. We related genes from various disease classes to function using Reactome or DAVID enrichment analysis tools [64, 101].

## Acknowledgements

We thank the Grishin lab members, Jing Zhang in particular, for helpful discussions. The study is supported in part by the grants (to NVG) from the National Institutes of Health (GM127390) and the Welch Foundation (I-1505).

