## supplemental Figures S1 and S2 for "Mutation severity spectrum of rare alleles in the human genome is predictive of disease type"

Supplemental Figure S1. Neural network structure.

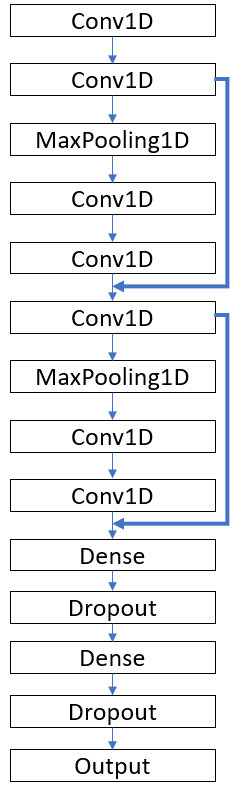

Supplemental Figure S2. ClistVis heatmap of potential disease-causing genes (UniProt label on the right) and genes with pathogenic SAVs (UniProt label on right with _P). Scores (labeled below) for each gene are colored from blue (low) to red (high) and clustered (20 clusters delimited by spaces) according to complete linkage of correlation distances. Two clusters with low AvgAI scores (mutation resistant) have a relatively high proportion of genes with known pathogenic variants and could help identify new disease-associated genes (red labels).

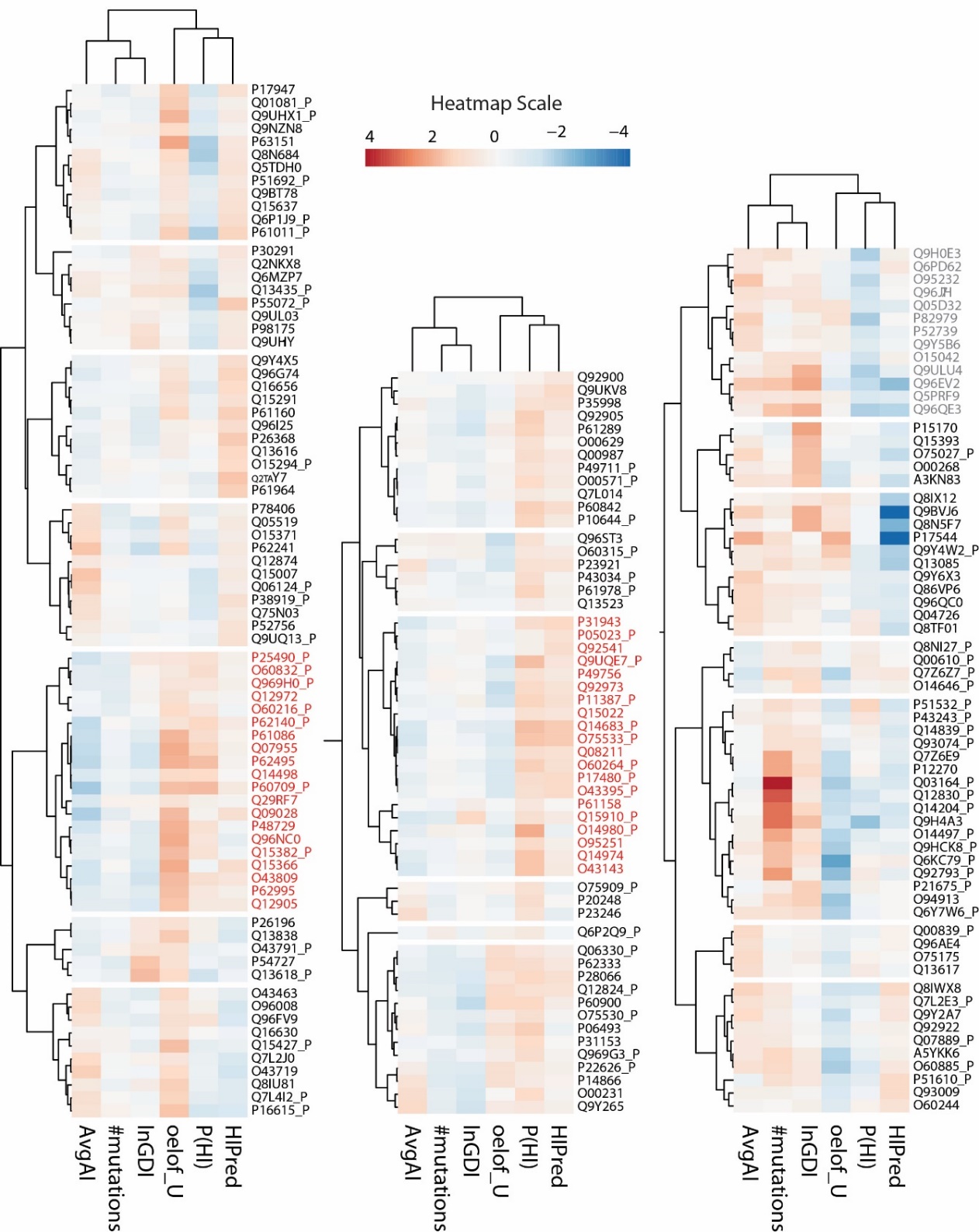
